# A conveyor feeder for animal experiments

**DOI:** 10.1101/801993

**Authors:** Jinook Oh

## Abstract

Several different types of open source feeders have been used in animal experiments in cognitive biology, neuroscience, psychology and related fields. These feeders use either dry pellets, which have hard surface and simple shape, or liquid food types such as sucrose solution. These food types can be rather easily manipulated due to its physical attributes. Although it is beneficial in terms of controllability, animal subjects often lose motivation to interact with operant conditioning devices offering such food items. Using natural food items such as fruits, insects, worms and pieces of meat will be helpful to keep the subject’s motivation high, however, those food items are not very well suited for currently available open source feeders due to its physical attributes, including its complex shape, sticky and delicate texture. We made a feeder to deliver such natural food items to animal subjects for operant conditioning, using relatively cheap and easily obtainable parts. For a validation, we built a full operant conditioning device for wolves and dogs, containing two of these feeders, a pressure-sensitive monitor and a speaker.

**Specifications table:** 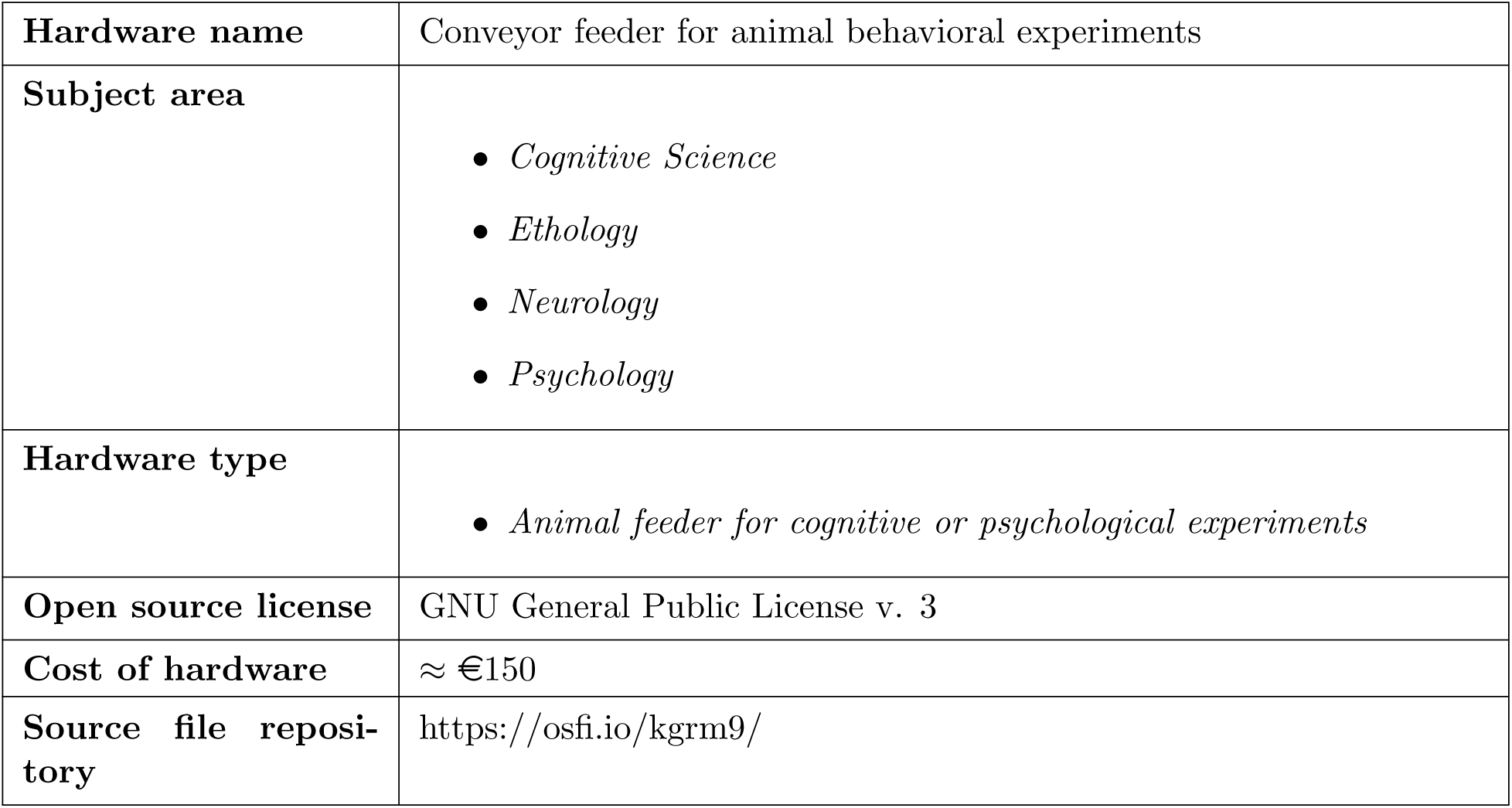

## 1. Hardware in context

Automated devices have been used for animal experiments in many scientific fields such as cognitive biology, neuroscience, ethology and psychology to test cognitive abilities and measure behaviors [1, 2, 3, 4, 5, 6, 7]. Also, open source devices have been increasingly used in these fields to facilitate reproduction, validation and customization [8, 9, 10].

An automatic feeder is a crucial part in such devices in animal experiments because it motivates animal subjects to participate for the experiments. Although several of them have been already developed [11, 12], feeders to release high quality food items is rare. Most automatic feeders for animal experiments use either a solid item with simple shape such as dry pellets and plant seeds, or a liquid item such as fruit juice and sucrose solution. These food items can be easily controlled by mechanical devices due to its physical attributes such as its regular shape, hardness, fluidity and so on.

A problem of using easily controllable food items is that it is rather common to see motivations of animal subjects decreasing (quickly or gradually depending on individuals) over time. This is especially true when the animal species on investigation is omnivorous and consume different food types in its natural diet. Such animal subjects include monkeys, dogs, wolves, cats, crows, ravens and parrots, and their natural diet includes meat, insect, worm, fruit, seed, root and so on.

When an animal cognition is investigated, many trials and sessions are required to conduct an experiment. Sometimes, a number of training sessions over a period of months is required just to reach a point to be ready for an actual experimental test. When motivation of an animal subject to the food reward is too low, the resultant data become rather unreliable because it becomes ambiguous whether the cognition of the subject is not capable of solving the given task or the subject simply lost its interest. Therefore, it is important to keep the subject animal’s motivation high during potentially long period of training and testing periods.

However, rather natural food items such as dead insects, worms, cut pieces of fruits, meat balls and sausages have delicate surface. Also, they sometimes have complex shapes. These food items can easily get caught by a gear or other moving parts of a mechanical device. Once it is stuck between certain moving parts, the food item is easily squashed and spread over different parts causing problems including compromised food rewards, malfunctioning of the device and so on.

In this paper, we describe a conveyor type feeder to help biologists for using natural food items to keep animal’s motivation high in training and experimental sessions. The feeder was designed with relatively cheap parts which are available in local hardware stores and online shops. We built the described feeder in an operant conditioning device composed of two conveyor feeders, a pressure-sensitive monitor and a loudspeaker to test dogs (*Canis lupus familiaris*) and wolves (*Canis lupus*). The operant conditioning device is briefly described in ‘Validation and characterization’ section.

## 2. Hardware description

The conveyor feeder is an open source automatic feeder which has many (33 in the current design) containers, each of which delivers a food reward to an animal subject in training or testing trials of psychological experiments. A 260 cm long timing belt is put on pulleys at top and bottom ends. Each food container is attached to this rubber belt on every 4 cm. All attached food containers take about half of the length of belt. At the beginning of a session, all the containers should be in upright positions so that they can be filled with food rewards by an experimenter. At the top end, the pulley is connected to a geared DC motor which is controlled by a motor driver, connected to an Arduino chip. At the bottom end, a piezoelectric sensor is positioned to sense a drop of a food piece. The sensor is also connected to the Arduino chip. When a signal to release a food reward is sent to the Arduino chip from an experimenter’s computer, the motor slowly rotates. As the motor, pulley, belt and containers move accordingly, one of the containers at the bottom side will go around the lower pulley and be flipped. Then a food piece in the flipped container will be dropped onto the piezoelectric sensor, sending a signal to the Arduino chip to stop the motor rotation.

Most open source feeders cannot deliver different types of natural food items, as described in the section 1. Although a conveyor type feeder was used in an operant conditioning device in [9], the feeder was limited on customization because it was built with several parts from a commercial toy. Using containers attached on a conveyor belt is a straight-forward idea and it has been used for long time, however, good sources of building details for scientific experiments with animal subjects are still scarce in open source hardware resources. What we try to offer with this report is providing one source of building experimental hardware for researchers in biological fields.

Our conveyor feeder

- enables using many types of quality food items for animal subjects
- can be built with commonly available parts, making further customizations feasible
- reduces cost of experimental device, when compared to commercial ones

## 3. Design files

### 3.1 Design Files Summary

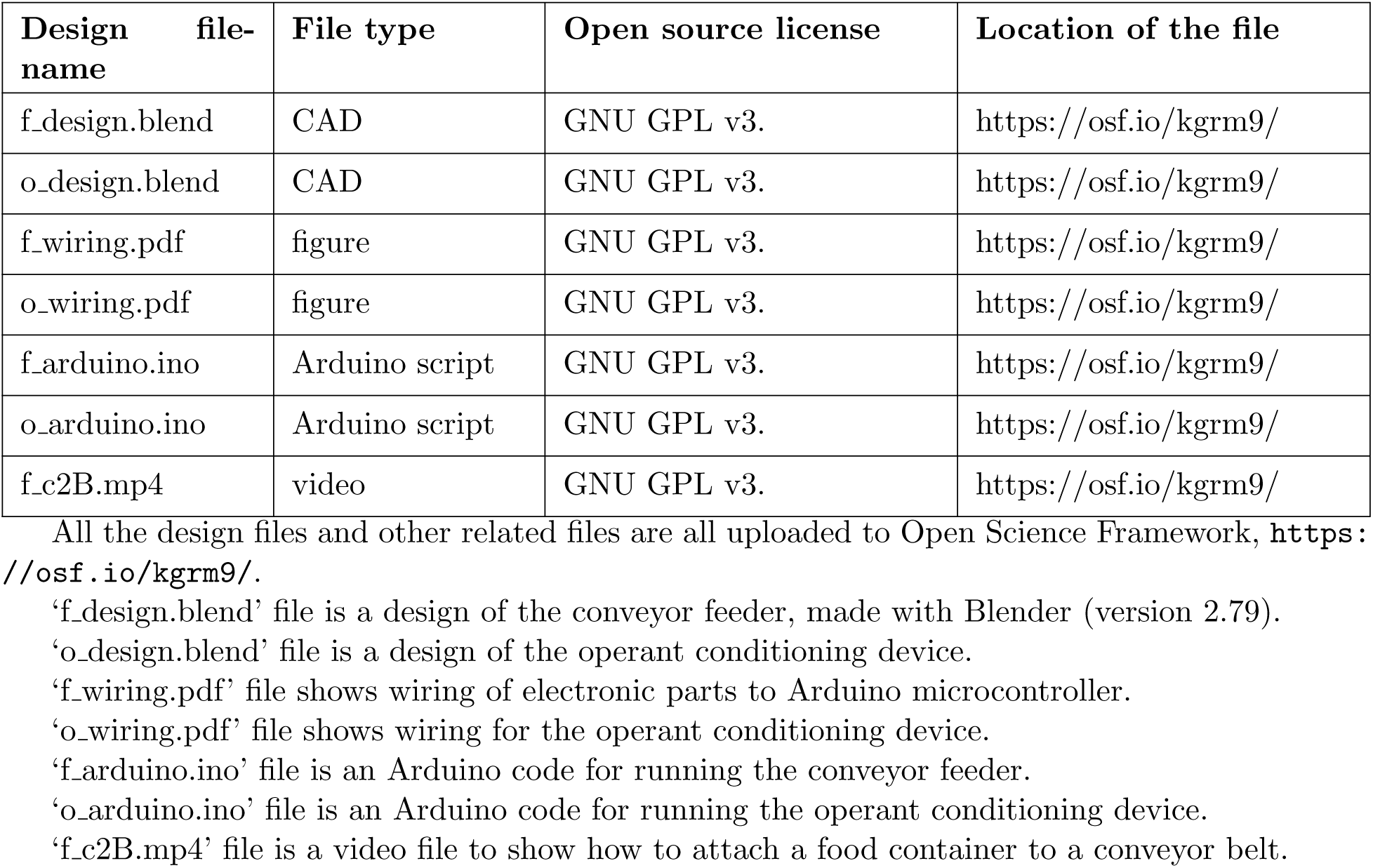

## 4. Bill of materials

| Designator | Component | Number | Cost per unit<br>currency (€) | Total cost (€) | Source of<br>materials |
| --- | --- | --- | --- | --- | --- |
| fe1 | Arduino UNO R3 | 1 | 20.07 | 20.07 | amazon.de |
| fe2 | 12V DC Power<br>supply adapter | 1 | 9.98 | 9.98 |  |
| fe3 | L298N motor driver | 1 | 5.79 | 5.79 |  |
| fe4 | 50 mm piezoelectric<br>sensor | 1 | 2.93 | 2.93 |  |
| fp1 | Acrylic clear tube<br>30/26/1000 mm | 1 | 6 | 6 |  |
| fp2 | Pack of 20 clear<br>acrylic sheet round<br>circle 1/8" | 2 | 5.81 | 11.62 |  |
| fe5 | Low RPM Gear<br>Motor | 1 | 32.81 | 32.81 | robotshop.com |
| fe6 | Mini push button<br>switch | 2 | 0.36 | 0.72 |  |
| fe7 | Resistor 1 M Ohm<br>(20pk) | 1 | 0.95 | 0.95 |  |
| fp4 | 6-32 x 1/4" Socket<br>Head Machine<br>Screw (25 pcs) | 1 | 1.51 | 1.51 |  |
| fp5 | 6mm D Shaft (10<br>cm long) | 1 | 6.34 | 6.34 |  |
| fp6 | Actobotics<br>Aluminum<br>Clamping Motor<br>Mount | 1 | 6.27 | 6.27 |  |
| fp7 | Flat Aluminium<br>Channel Bracket | 1 | 1.15 | 1.15 |  |
| fp8 | MakeBlock 3m<br>Open-end Timing<br>Belt | 1 | 10.25 | 10.25 |  |
| fp9 | MXL 60 Tooth<br>Timing Pulley | 2 | 2.03 | 4.06 |  |
| fp10 | Lynxmotion Pillow<br>Block - 6mm PB-02 | 2 | 7.65 | 15.3 |  |
| fp11 | Metal rod (3 mm<br>diameter, 6 cm<br>long) | 1 | 3 | 3 | Bauhaus<br>(local hardware<br>store) |
| fp12 | Wooden panel (20<br>x 15 x 1.2 cm) | 1 | 10 | 10 |  |

## 5. Build instructions

To build the feeder, we used the tools shown in Table 1.

**Table 1:**
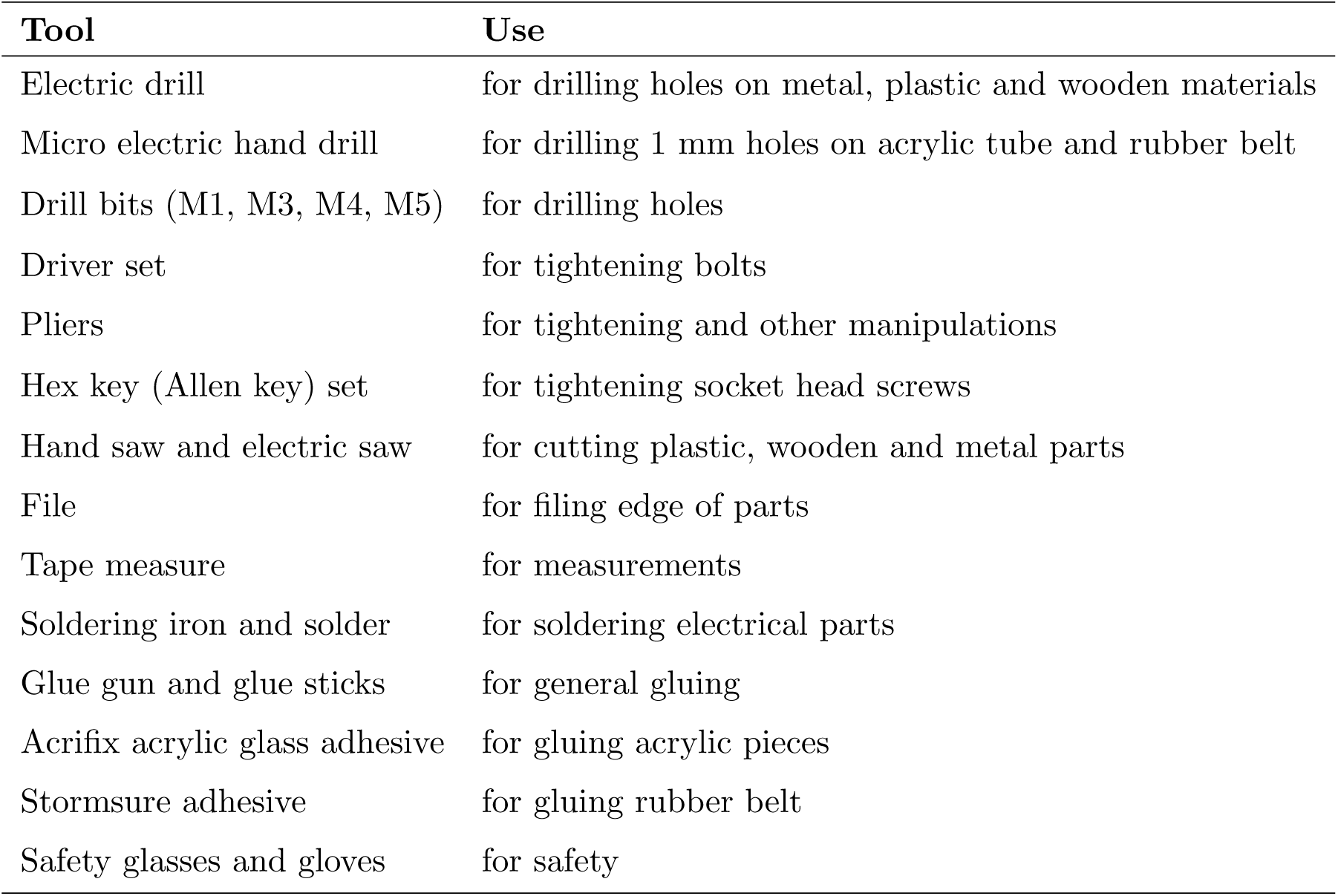
Used tools to build the feeder

| **Tool** | **Use** |
| --- | --- |
| Electric drill | for drilling holes on metal, plastic and wooden materials |
| Micro electric hand drill | for drilling 1 mm holes on acrylic tube and rubber belt |
| Drill bits (M1, M3, M4, M5) | for drilling holes |
| Driver set | for tightening bolts |
| Pliers | for tightening and other manipulations |
| Hex key (Allen key) set | for tightening socket head screws |
| Hand saw and electric saw | for cutting plastic, wooden and metal parts |
| File | for filing edge of parts |
| Tape measure | for measurements |
| Soldering iron and solder | for soldering electrical parts |
| Glue gun and glue sticks | for general gluing |
| Acrifix acrylic glass adhesive | for gluing acrylic pieces |
| Stormsure adhesive | for gluing rubber belt |
| Safety glasses and gloves | for safety |

### 5.1 Building frame

Our feeder was tested in an operant conditioning device composed of two conveyor feeders, a pressure-sensitive monitor and a loudspeaker. The procedure to build a frame to contain all these components will be described here only briefly, because the focus of this paper is on the feeder and frames in other applications could be completely different in its shape and material.

Metal beams (2 cm thick) and corresponding connecting parts (triangular corner pieces, bolts and sliding pieces for screwing bolts in) were purchased from Haberkorn (haberkorn.com) and assembled using a hex key as shown in Figure 1. Three parts were assembled first as shown in the 1st, 2nd and 3rd image of Figure 1). Four long (1.4 m) beams were connected on top of the assembled part (1) as shown in the 4th image of Figure 1. The 5th image of Figure 1 is the finished frame, in which the part (2) and (3) were attached to the middle and top side of the 1.4 m long beam. Also, it shows two 1.3 m long beams, attached between the assembled part (1) and (3). Motors of two conveyor feeders were positioned on top of the part (3) above those beams. Parts of a feeder was aligned along this 1.3 m long beam.

**Figure 1:**
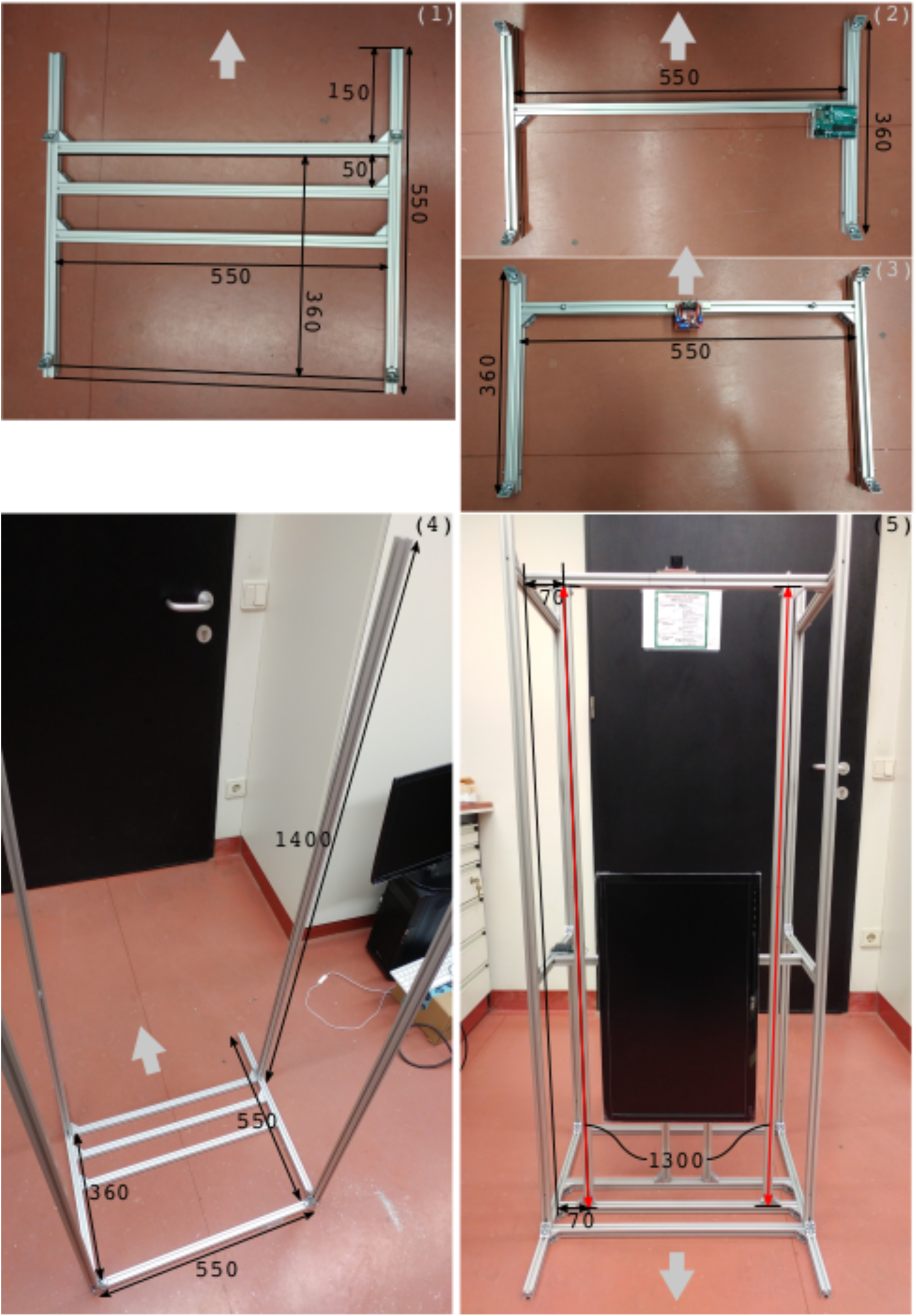
Building frame of the operant conditioning device. Numbers are in millimeters. Grey color short arrows indicate the front side of the device.

### 5.2 Preparing belt with food containers

The acrylic clear tube (fp1) was cut into smaller pieces as shown in the 1st image of Figure 2. The length of each cut tube was approximately 18 mm. To make a bottom of a food container, one of clear acrylic round plates (fp2) was glued to a cut tube, using Acrifix adhesive. The glued pieces were dried under sunlight for a half hour (the 2nd image of Figure 2). Thirty three containers were prepared. Running thirty trials in one experimental session was planned, but three additional containers were attached just for cases of erroneous behaviors such as a failure of sensing a food drop.

**Figure 2:**
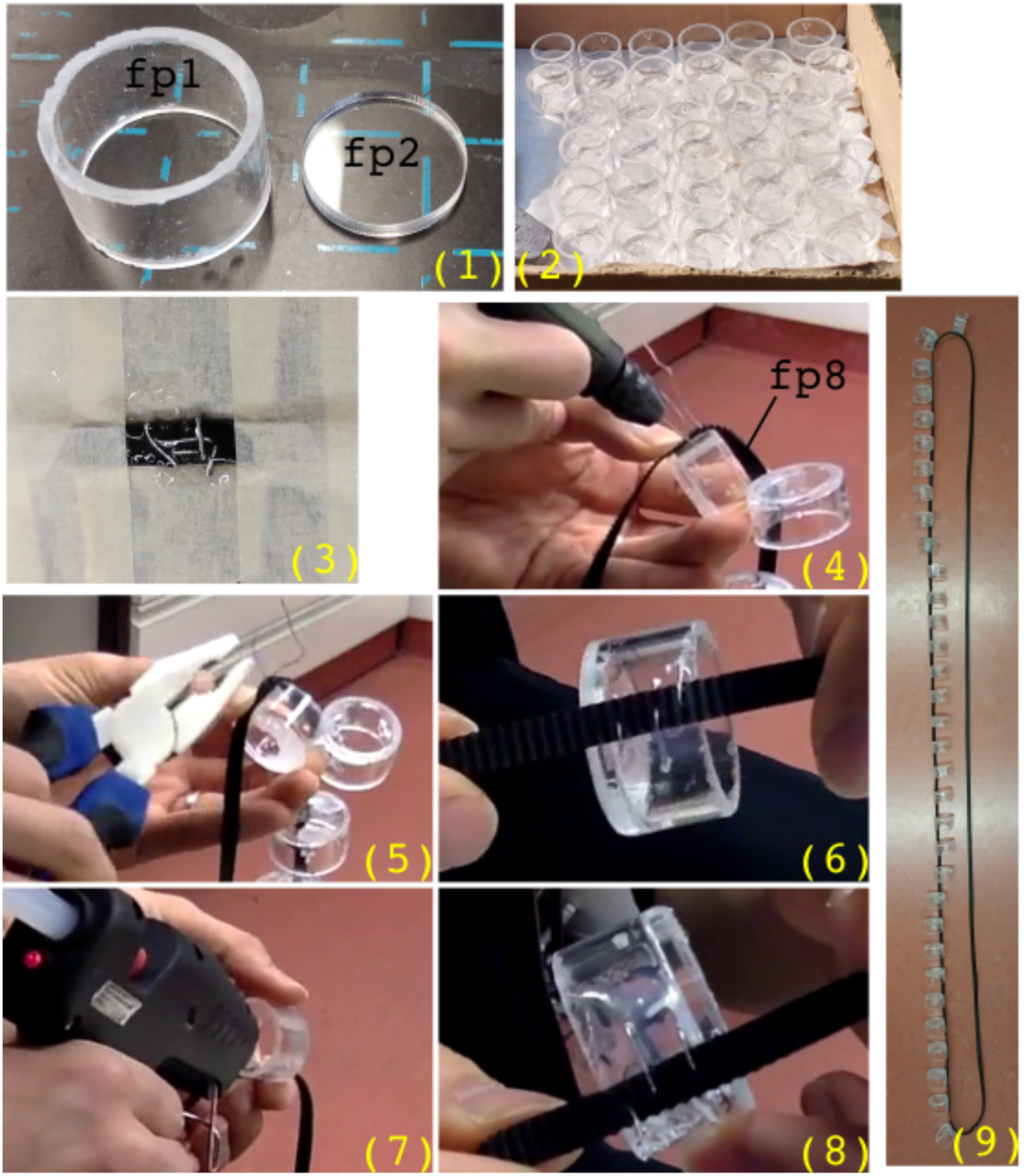
Preparing belt with food containers

A timing belt (fp8) was cut to 260 cm long, then its ends were attached to each other to form a conveyor belt to turn around pulleys. To attach both ends together, several holes were drilled around both ends of the belt using a micro electric hand drill, then a plier and thin (0.6 mm) wire were used to sew the ends through the holes. Extra adhesive (Stormsure) was used to fortify the attachment. Drying applied adhesive after sewing with a wire is shown in the 3rd image of Figure 2.

The belt and a container were sewed together with the thin wire through several holes as shown in the 4th and 5th image of Figure 2. The sewn container looked like the 6th image of Figure 2. The attachment was quite robust already before gluing. The glue gun was used to fortify the attachment and cover sharp ends of the wire (the 7th and 8th image of Figure 2). The process of this attachment, (4-8), is also shown in a video, ‘f_c2B.mp4’ uploaded to https://osfi.io/kgrm9/. The distance between two containers (from top of a container to top of the next container) was approximately four centimetres. The 9th image of Figure 2 shows the belt with food containers after all the preparations.

### 5.3 Building upper part

The conveyor feeder here is vertically aligned. The upper part has a motor (fe5) to rotate a pulley (fp9) to move a timing belt (fp8). A clamping style motor mount (fp6) was fixed on a flat bracket (fp7) using socket head machine screws (fp4). After fixing the bracket on top of the frame, the motor was clamped in the motor mount with a hex hey. The pulley was attached to the motor, also using a hex key. Finally, a motor driver (fe3) was positioned near to the motor. The finished part is shown in Figure 3.

**Figure 3:**
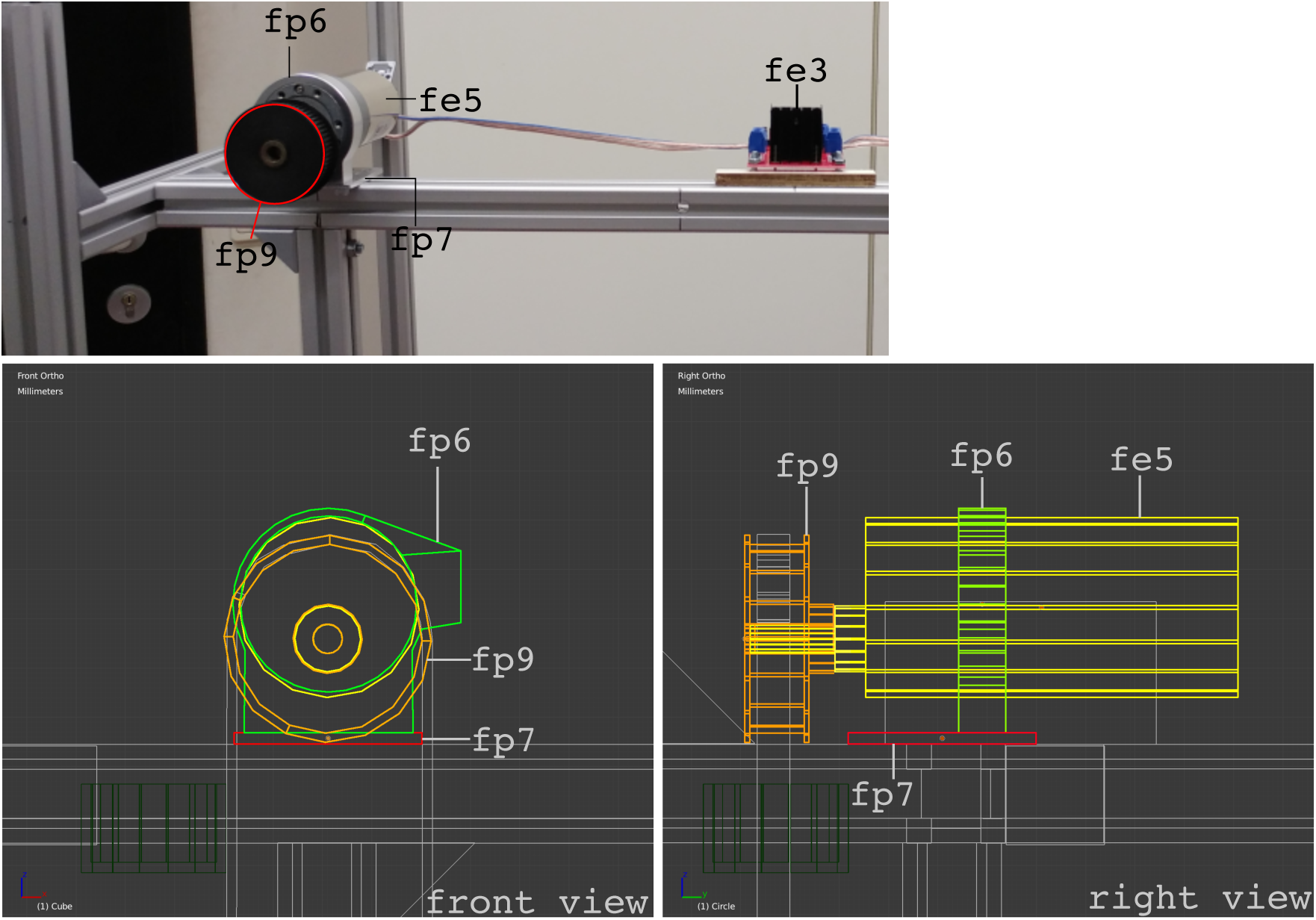
Upper part of the feeder.

### 5.4 Building lower part

The lower part of the feeder is consisted of another pulley, pillow blocks to keep the pulley with a D-shaft in position and a piezoelectric sensor to detect a drop of food piece. Two pillow blocks (fp10) were attached, using screw nails, to wooden pieces, cut from a wooden panel (fp12). The attached pieces are shown in the middle image of Figure 4. Two sides of a wooden piece were filed, as shown in the lower image of Figure 4, to allow smooth passing of food containers. A metal D-shaft (fp5) was cut to a 5 cm long piece. The cut shaft was inserted to a pulley, then both ends of the shaft were inserted to pillow blocks. The prepared belt with containers from section 5.2 was put on the upper side pulley and the lower side pulley. Then the wooden pieces with pillow blocks were attached to two metal beams as shown in Figure 5. Finally, the position of the lower pulley on the D-shaft was adjusted to make the upper side pulley, the belt and the lower side pulley are all aligned along a straight vertical line.

**Figure 4:**
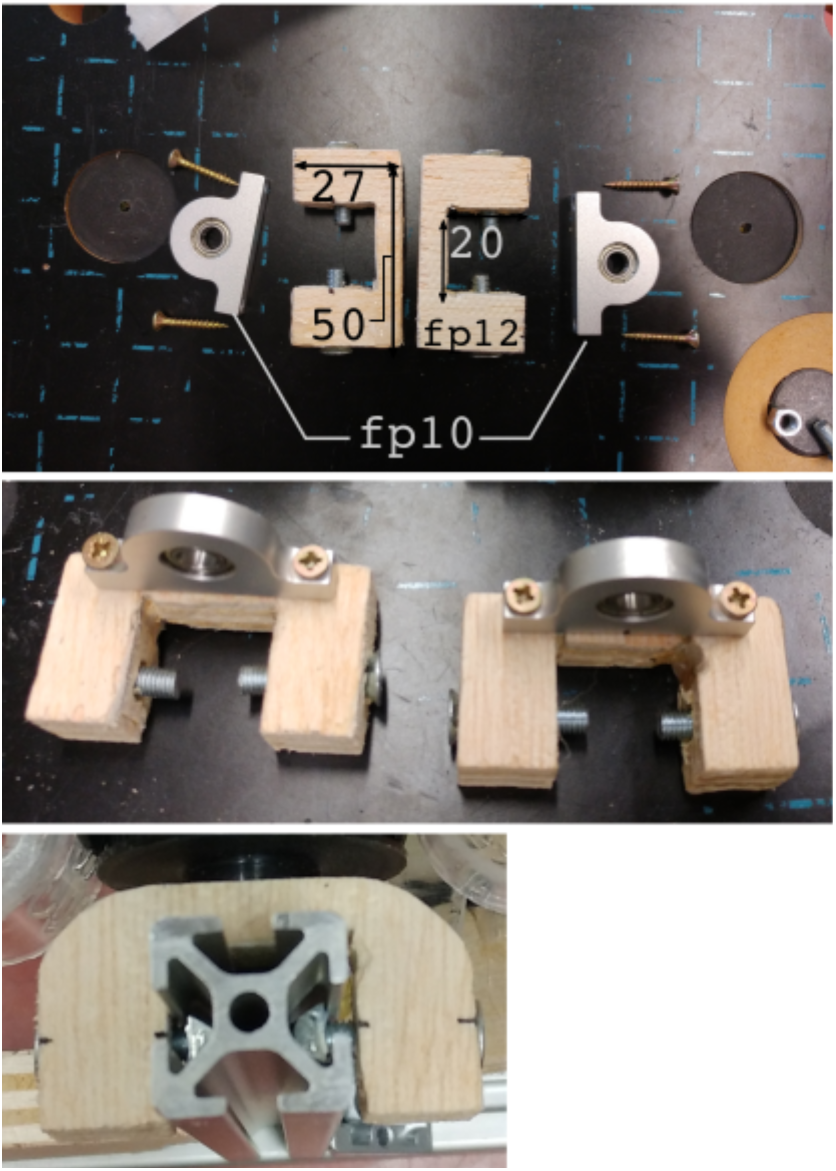
Preparing two pillow blocks. Numbers are in millimeters.

**Figure 5:**
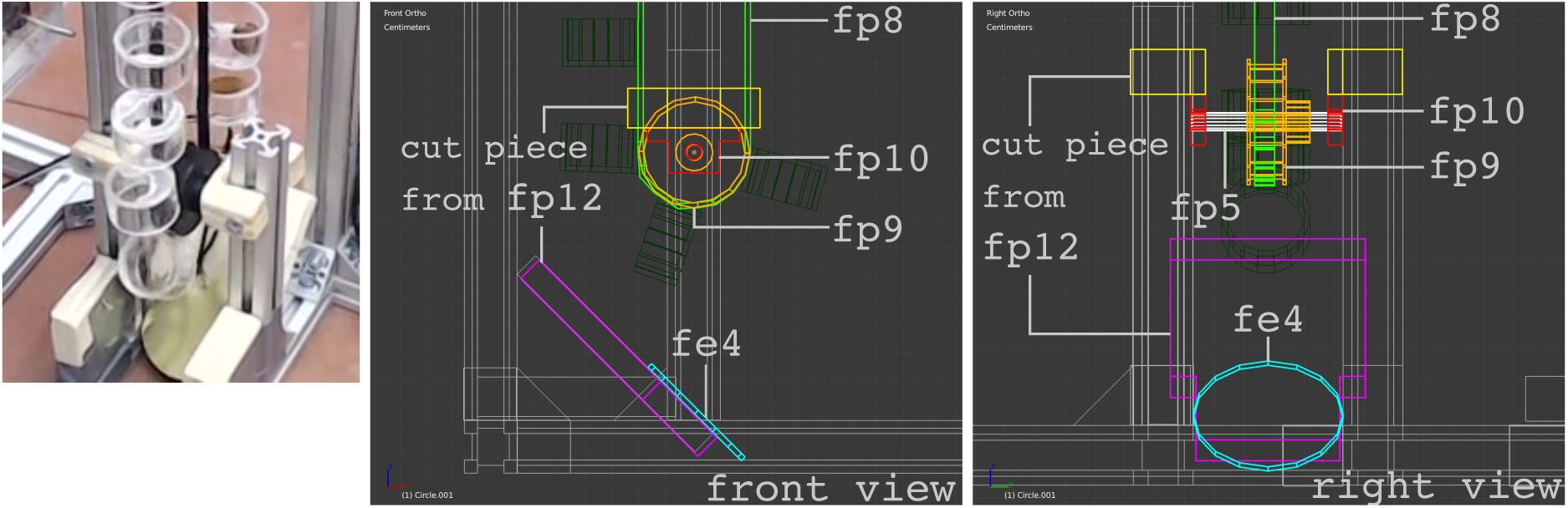
Lower part

A wooden piece, shown in Figure 6, to support the piezoelectric sensor was cut from the wooden panel (fp12), then two holes were drilled to receive the 3 mm thick metal rods (fp11). A piezoelectric sensor (fe4) was soldered with cables and attached to a wooden cut piece (fp12) with a glue gun. The metal rod (fp11) was cut to two 3 cm long pieces. By inserting the cut rods into holes in triangular corner pieces (purchased with metal beams from Haberkorn) and the wooden piece, the piezoelectric sensor was positioned below the lower side pulley to make a falling food piece to hit the sensor. (See Figure 6 and 7, see also Figure 5 for the finished lower part)

**Figure 6:**
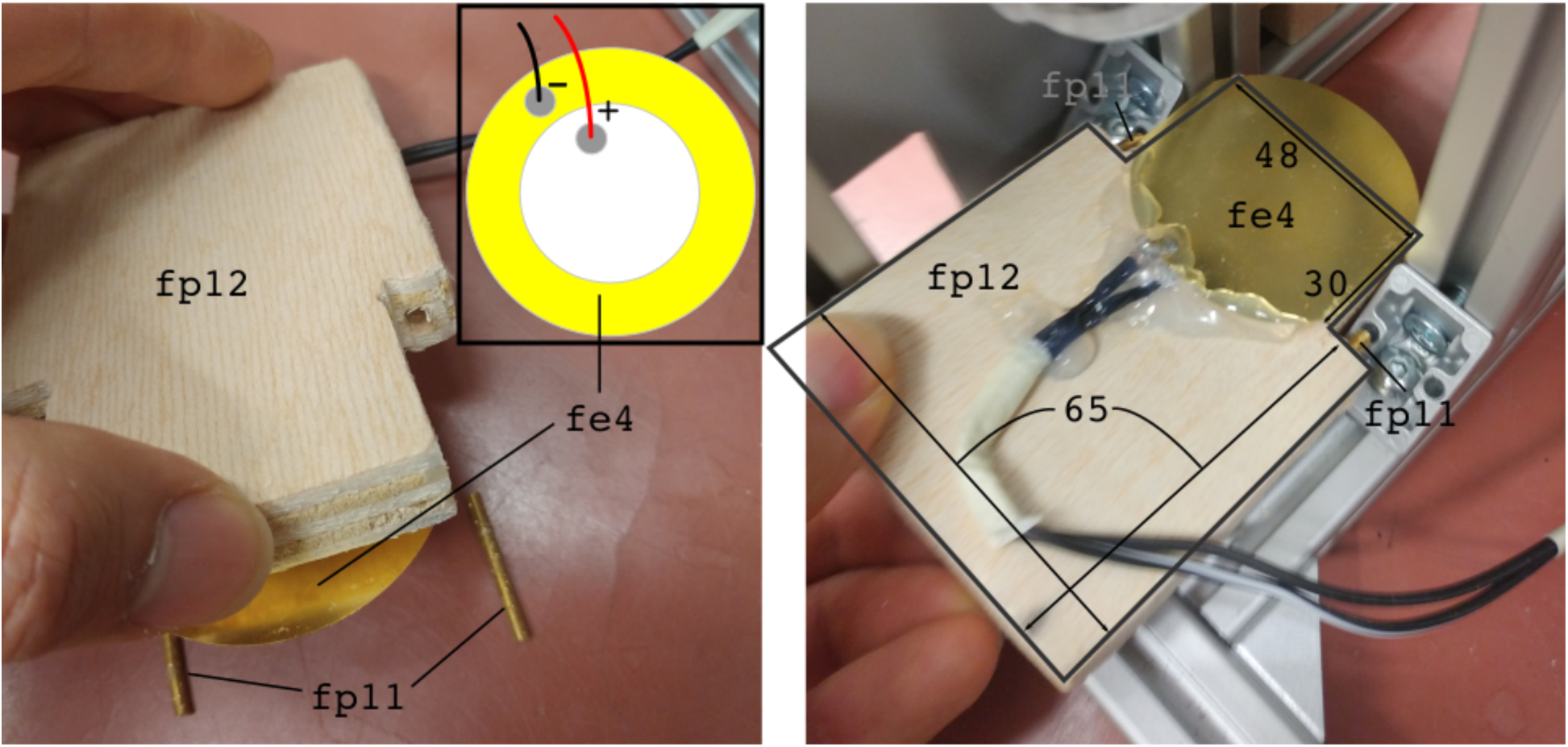
Positioning piezoelectric sensor. Numbers are in millimeters.

**Figure 7:**
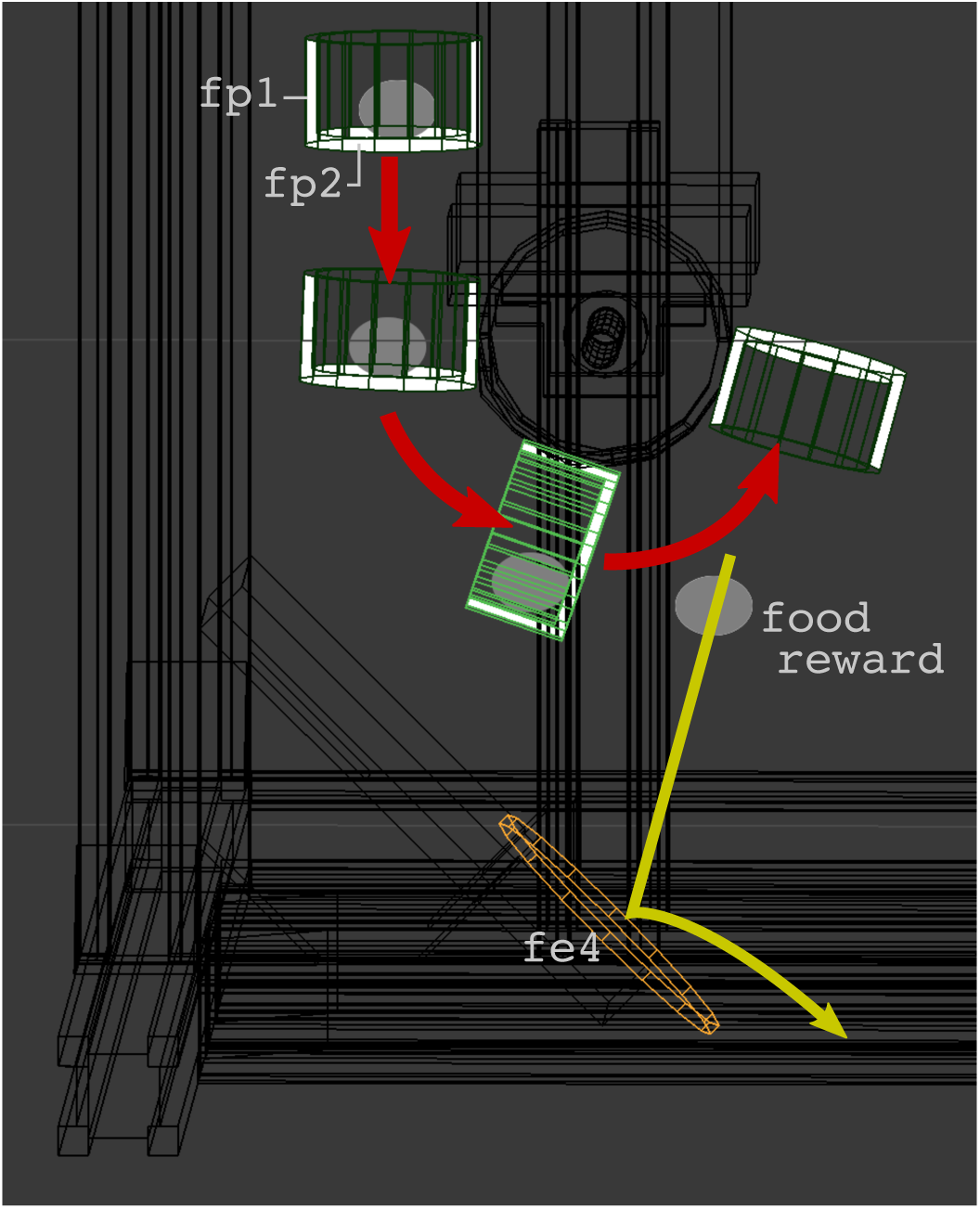
Releasing a food reward

### 5.5 Wiring

The wiring diagram is shown in Figure 8. A motor (fe5) was connected to a L298N motor driver (fe3) which was connected to an Arduino microcontroller (fe1) and powered by a 12V DC power supply (fe2). A piezo-electric sensor (fe4) was connected to an analog pin on Arduino chip with a 1 MOhm resistor. Additionally, two mini push buttons were connected to Arduino chip for cases when manual and continuous motor running is necessary. For example, when all trials are finished, all food containers are flipped and positioned in an opposite side from its original position. An experimenter can keep rotating the motor with the push button to position all containers in its original position to refill the containers.

**Figure 8:**
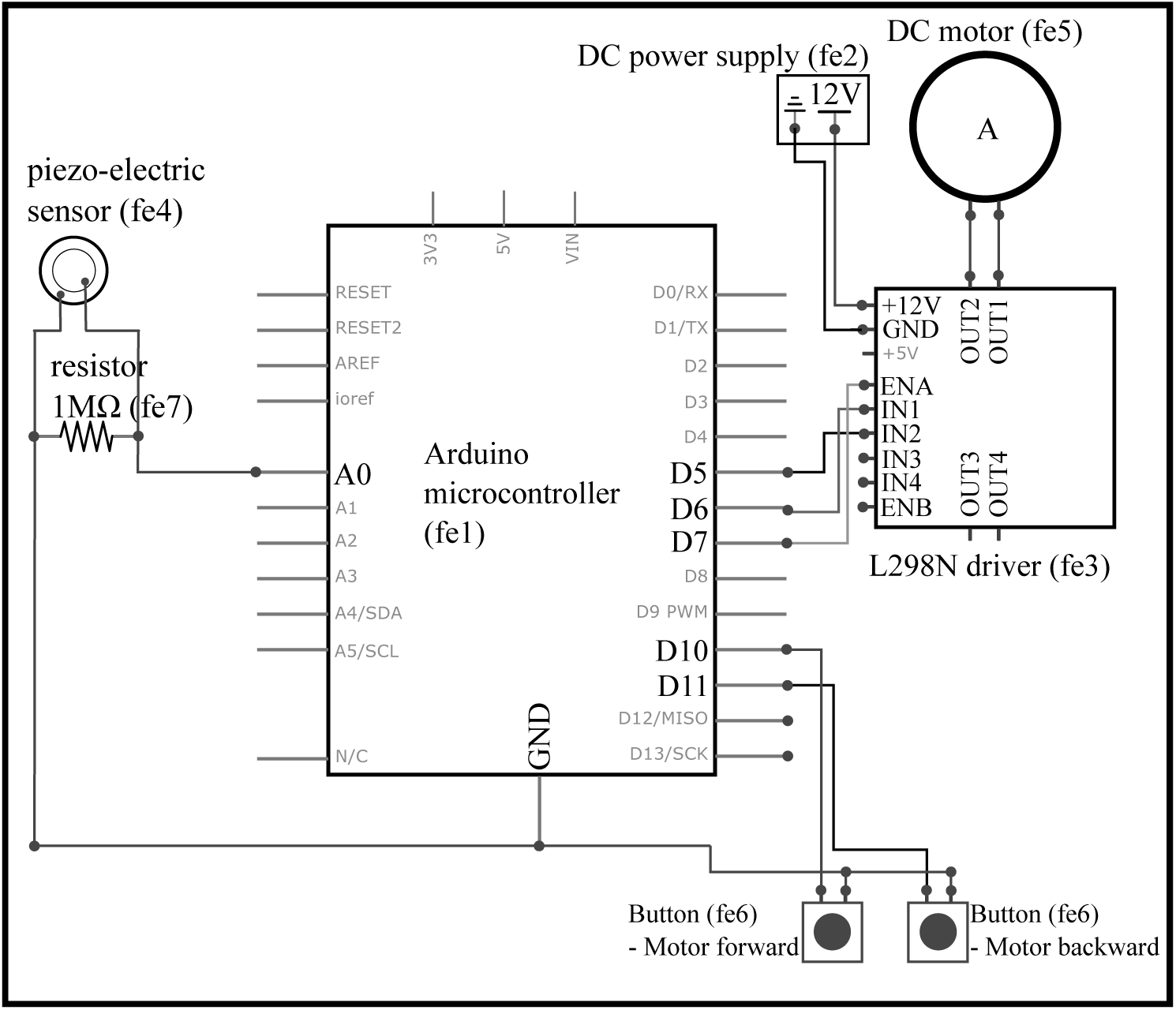
Wiring electronic parts to Arduino chip

### 5.6 Software

All the software developed during this project was uploaded to https://osf.io/kgrm9/, and can be freely downloaded.

## 6. Operation instructions

One can test the conveyor feeder using the test script, test.py on https://osf.io/kgrm9/. Following steps should be conducted to test the feeder.

a. Download Arduino software from https://www.arduino.cc/ and install it.
b. Download Python (Python 2 or 3) from https://www.python.org then install it. If the operating system is Windows, add Python installation path and ‘[PythonInstallationPath]*\*Scripts’ folder to ‘Environment variables’.
c. Open a terminal window and type ‘pip install pyserial’ to install PySerial library to communicate with Arudino microcontroller.
d. Load ‘f_arduino.ino’ file in Arduino software.
e. Connect Arduino chip to a computer using a USB cable.
f. Select a board (Tools >Board >Arduino/Genuino UNO).
g. Select a port (Tools >Port >[port that appears when Arduino UNO is connected to computer]).
h. Press ‘Upload’ button to upload it to Arduino chip.
i. Power the motor driver (fe3) with 12V DC power supply.
j. Run ‘test.py’ with a command, ‘python test.py’, on a terminal window or using other apps such as IDLE (included software in Python package).
k. Type ‘test’ and enter.

## 7. Validation and characterization

Two conveyor feeders were built in an operant conditioning device (Figure 9) to test dogs and wolves. The wiring diagram for this implementation is shown in Figure 10.

**Figure 9:**
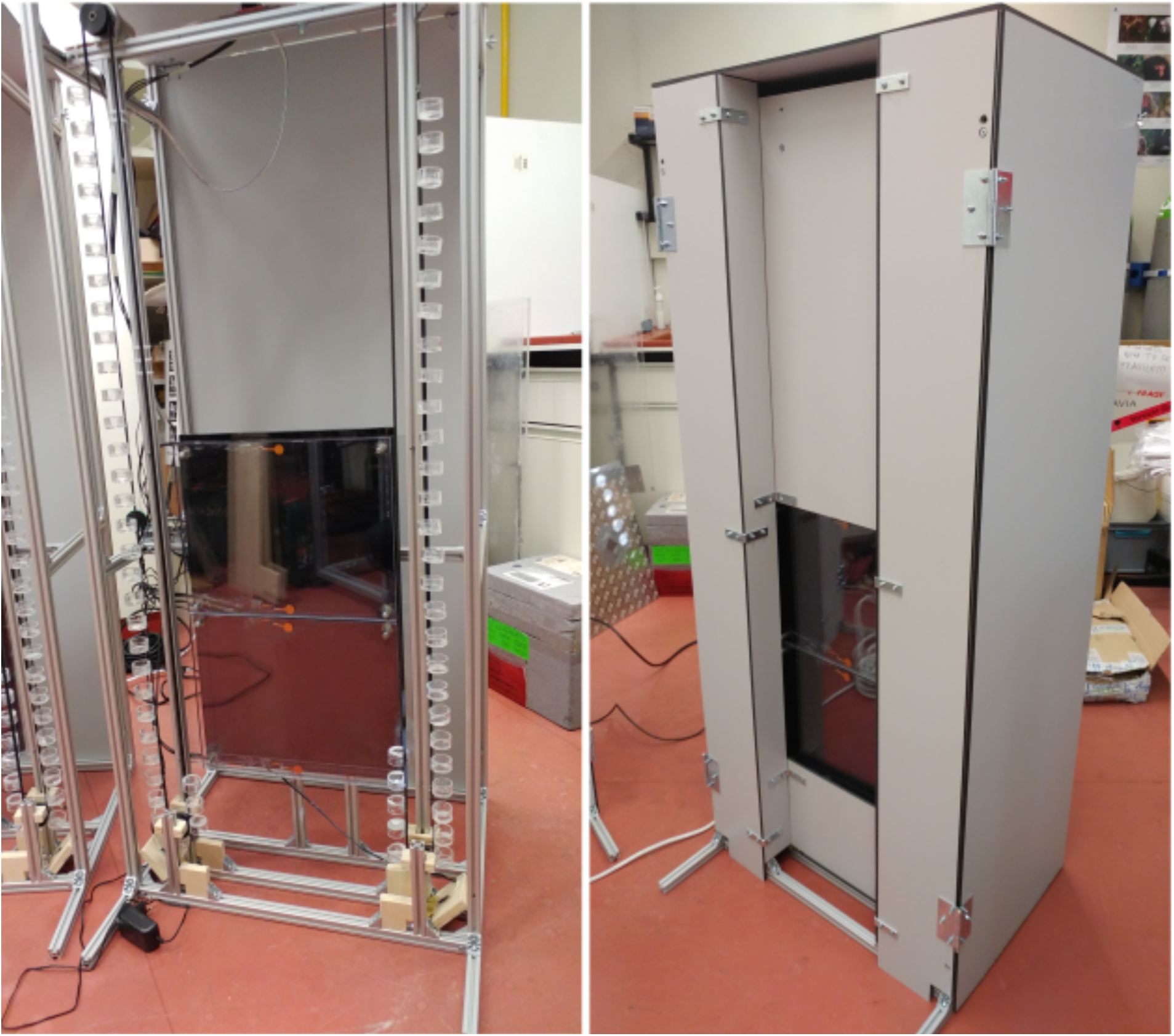
Testing operant conditioning device

**Figure 10:**
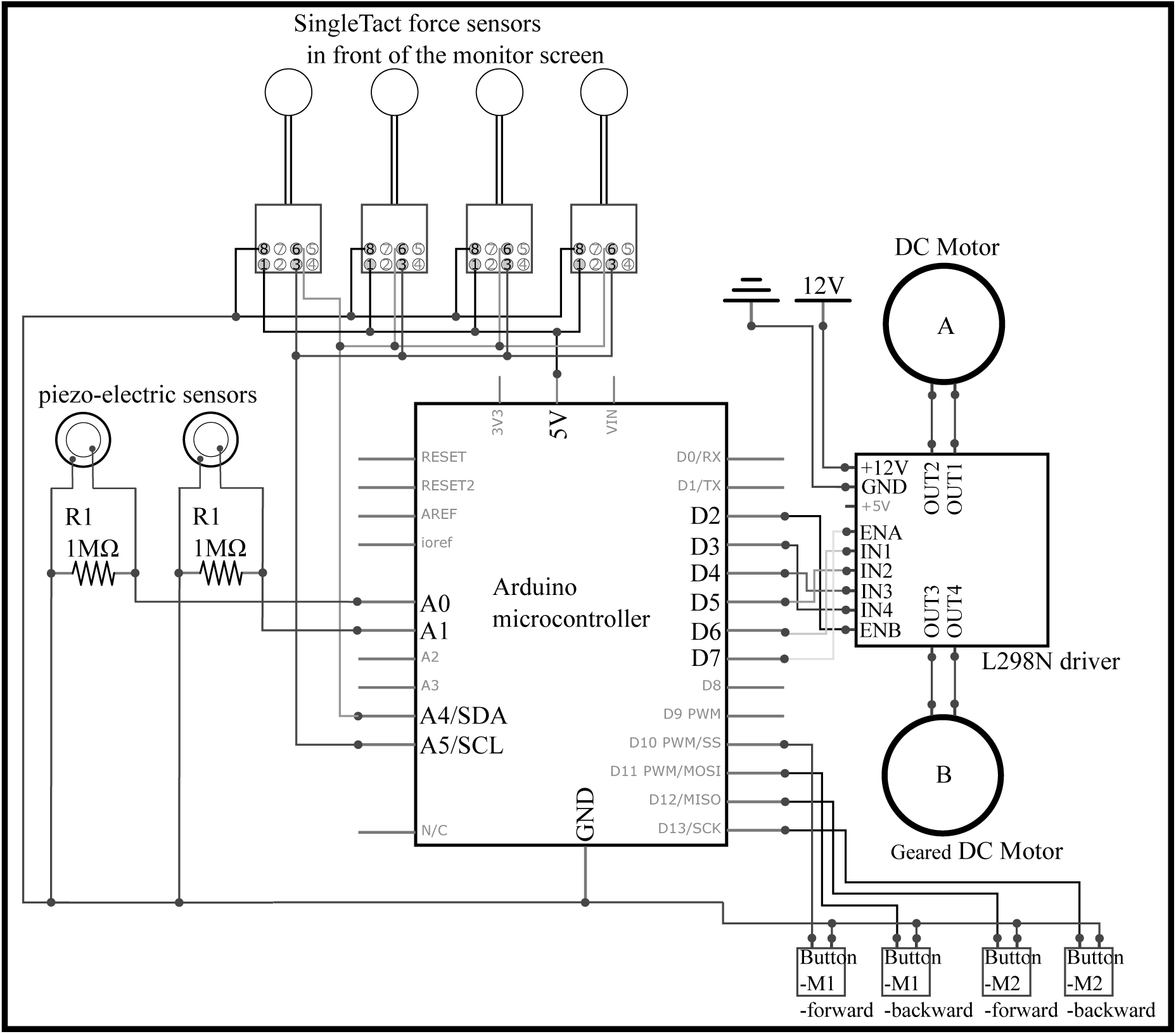
Wiring for the operant conditioning device

Two layers of transparent acrylic plates were placed in front of a monitor and four force sensors were placed between the layers to detect strong enough press of a subject on a stimulus image. We implemented this force-sensitive plates instead of using a commercial touchscreen due to two reasons. First, a touchscreen (capacitive or infrared type) can show malfunctioning depending on an individual dog or wolf due to the subject’s excessive saliva on touchscreen. Saliva can be excessively produced with expectation of feeding and dropped on screen, then it can cause false touch recognitions. Second, force sensors can be used to detect a meaningful press of an animal subject, while a commercial touch screen often detects an unintentional touch with body or tail of the subject.

Each feeder was filled with dry pellets and cut sausages respectively to provide low and high quality food rewards. In testing, two images were displayed on upper and lower half of the screen. Depending on the subject’s choice, one of the two feeders was activated to release a food reward.

The feeders were first tested for over 300 food releases using cut sausages and almond seeds. No noticeable error was found during this pre-test. Then, the feeder was tested with a wolf for 90 trials. During this test, two erroneous behaviors of the feeder occurred. When the motor just started to rotate to release a food reward, the subject hit the device with its body, shaking the entire device. The piezoelectric sensor picked up the vibration from the shaking and stopped the rotation of motor before a release of a food piece. This error could be eliminated by installing an extra protective layer in front of the device.

Also, the piezoelectric sensor can fail to detect a food drop. This error did not occur during our tests, however, it is possible in the case of using very light-weight and soft food piece. To prevent this error, the threshold to detect a food drop in Arduino code can be lowered. If the error persists, one can use a different sensor such as a break beam sensor or piezo vibration sensor (film type with mass).

## 9. Human and animal rights

The housing conditions and the experimental design were in accordance with Austrian legislation and the European Association of Zoos and Aquaria (EAZA) husbandry guidelines.

